# Mapping the auditory space of *Culex pipiens* female mosquito in 3D

**DOI:** 10.1101/2023.01.09.523250

**Authors:** Dmitry. N. Lapshin, Dmitry. D. Vorontsov

**Affiliations:** Institute for Information Transmission Problems of the Russian Academy of Sciences (Kharkevich Institute) Bolshoy Karetny per. 19, Moscow, 127994, Russia; Koltzov Institute of Developmental Biology Russian Academy of Sciences Vavilova 26, Moscow, 119334, Russia

## Abstract

The task of directional hearing faces most of the animals that possess ears. They approach this task in different ways, but the common trait is the usage of the binaural cues to find the direction to the source of sound. In insects, the task is further complicated by their small size and, hence, minute temporal and level differences between two ears. A way to overcome this problem is to receive the particle velocity component of sound rather than the pressure, as the former naturally involves directionality. However, even in this case, one ear is not enough for directional hearing: a single symmetric flagellar particle velocity receiver cannot discriminate between the two opposite directions along the vector of the sound wave. Insects that use flagellar auditory organs, and mosquitoes in particular, possess a pair of receivers, which presumes the usage of binaural hearing. Its mechanisms are expected to be significantly different from the ones typical for the pressure receivers. However, the directionality of flagellar auditory organs has received little attention. Here we measured the in-flight orientation of a female mosquito antennae and obtained detailed physiological mapping of the Johnston’s organ directionality at the level of individual sensory units. By combining these data, we provided a three-dimensional model of the mosquito’s auditory space. The natural orientation of the antennae together with angular distribution of sensory units in each of the Johnston’s organs was found to be optimal for binaural hearing focused primarily in front of, above and below a flying mosquito.

## 1. Introduction

Sound localization is one of the major tasks of the auditory system. In animals, it is differently solved depending on which of the components of the sound wave, pressure or particle velocity, is being detected. In the systems based on pressure receivers, such as human ears, binaural cues like time- and level-differences are used to localise sounds (Schnupp and Carr, 2009). However, at low frequencies (below about 10 kHz for most animals) the amplitude difference at the two ears, as well as the timing difference may be very small even in mammals (Köppl, 2009), not to mention tinier creatures like insects. To solve this problem, two ears become mechanically or acoustically connected, thus creating pressure-difference receiver, which enables amplification of tiny acoustic cues into larger interaural differences which can be effectively processed by the nervous system (Miles et al., 1995; Michelsen and Larsen, 2007; Römer, 2015).

On the other hand, particle velocity receivers, such as the antennae of mosquitoes, are inherently directional (Belton, 1974; Robert, 2005; Morley et al., 2012). The radially symmetrical structure of the Johnston’s organ (JO), a modified second segment of insect antenna, implies that only a fraction of sensory cells that is aligned to the vector of a given acoustic wave will generate a significant response to it. As the air particles move back and forth during the propagation of a sound wave, a single mosquito JO and other similarly designed auditory receivers should not be able to distinguish sounds that come from the two opposite directions. As sound fields are usually strongly divergent close to small sound sources, bilateral particle velocity receivers such as the mosquito’s antennae may experience vastly different vector fields depending on the distance from the sound source, which may affect their ability to extract directionality data from the sound, i.e., the velocity field may or may not directly point to the sound source (Robert, 2005). For the sake of simplicity, one may assume that each antenna is functionally omnidirectional, and only the inter-antennal amplitude differences (IADs) based on the nonlinearities that produce difference frequencies are used to provide the localization cues (Ziemer et al., 2022). Although the modelling experiments in that study demonstrated that IADs in mosquito may be sufficient for the task, we consider it quite unnatural that the information on directionality that is mapped in the activity of the numerous auditory units of the JO may be lost on its way to the brain, even if it does not allow to unequivocally discriminate between the two opposite directions.

It is intuitively clear that paired antennae of an insect constitute a single sensory system, and that the synthesis of information from the two receivers expands the resolution of the whole auditory space. The most obvious approach to choose the correct direction to the sound source from the two opposite ones would be to compare the responses originating in the left and the right JOs based on the directional mapping provided by each of them. Insects rarely hold two antennae in parallel, thus the directional characteristics of individual sensory units belonging to the left and right JO’s overlap in some non-trivial way, possibly providing additional interaural cues for sound localization. The hypothetical presence of irregularities in the directional diagrams of each JO, mirrored between the left and the right, would speak in favour of the presence of a mechanism for binaural directionality.

Such irregularities were indeed demonstrated in the *Culex* male mosquitoes, with the two of the four quadrants of the JO being more populated with responding cells (Lapshin and Vorontsov, 2019). That result, however, may reflect not only the true functional asymmetry of the mosquito JO, but also the possible experimental bias of recording the auditory responses from only a fraction of the antennal nerve and, thus, from the units unevenly distributed around the axis of the antenna. The latter assumption could not be avoided, as the recording electrode in that study was inserted through the cuticle, the recording sites in the antennal nerve were selected blindly, based only on the level of auditory response, while the electrode could be directed towards the mosquito from a very limited angular range in an attempt to avoid touching the large extended antennae of a male mosquito.Additionally, due to the same spatial limitations of the experimental setup, the recording in that study was made unilaterally from only the left JO.

Some of the above limitations could be overcome by recording from the JO of a female mosquito. The antennae of a female are smaller than those of a male. In experiment, they allow more freedom of inserting the recording electrode, as well as to record from either the left or right JO. Variability of the electrode insertion position would significantly decrease the possible bias caused by blind selection of recording sites within the antennal nerve.

The question ‘What do female mosquitoes hear?’ demands much attention by itself. The auditory system of biting female mosquitoes is poorly studied compared to that of the conspecific males, the outstanding listeners. Our very limited knowledge on such a practically important subject can be at least partially explained by the fact that for a long time it was difficult to demonstrate any behavioural auditory responses of female mosquitoes, although they also possess a rather sophisticated auditory system.

When the effect of mutual flight tone matching between a male and a female was discovered (Gibson and Russell, 2006) it became possible to use it as an auditory response. When the flight tones of a male and a female are too different for 1:1 matching, which is normal for many mosquito species, they were found to converge at a higher harmonic (Cator et al., 2009). Since the male’s flight tone in many species is outside the frequency range of a female mosquito, as well as their common higher harmonic tone, both male and female mosquitoes detect difference tones produced by nonlinear mixing of their flight tones in the vibrations of the antennae (Warren et al., 2009; Pennetier et al., 2010; Simões et al., 2016; Simões et al., 2018), while the effect of harmonic convergence appears only as an unintended consequence of difference tone detection. The very existence and the nature of the auditory interactions in mosquitoes proved to be the points of contradiction: statistical analysis of flight tones in pairs of mosquitoes (Aldersley et al., 2016) demonstrated that harmonic convergence exists and that it is an active phenomenon, which does not occur by chance. Simões et al. (2016) proposed that harmonic convergence might be just an epiphenomenon – the unintended consequence of adjustments in the fundamental flight tones so that the resulting difference tones fall within the optimal frequency ranges for detection. A recent study of Somers (2022) showed that harmonic convergence could be only a random by-product of the mosquitoes’ flight tone variance and may not be a signature of acoustic interaction between males and females. In any case, the hearing frequency range of a flying female mosquito (Lapshin, 2013) allows it to detect a flight tone of a male in the form of the difference tone(s), which can serve a physiological basis for acoustic interaction in mating behaviour.

Another behavioural response to sound, the increase of flight speed in free-flying mosquitoes, may be reliable enough to measure the auditory thresholds in males (Lapshin and Vorontsov, 2021; Dou et al., 2021), while the auditory thresholds detected in the latter study for females of *Aedes aegypti* were very high (ca. 70 dB).

At the same time, there are examples of distant attraction to the communication sounds of frogs in blood-sucking midges (Toma et al., 2005; Borkent and Belton, 2006; Borkent, 2008) and mosquitoes (Bartlett-Healy et al., 2008; see the paper of Steele and McDermott, 2022 for review). A study by Menda et al. (2019) suggests that the *Aedes* female mosquitoes can be attracted to the sound frequencies similar to those of human speech. Although the results on the female mosquito audition are far from conclusive, from the existing behavioural studies one can reason that biting females of some dipterans, and mosquitoes in particular, can perceive the direction towards the source of a sound.

The lack of direct physiological data on auditory directionality in female mosquitoes, together with the above-mentioned experimental convenience, formed the basis of this study. To measure the directionality of the JO sensory units, we rotated the vector of acoustic waves around the antenna of a female mosquito while recording from the axons of the auditory neurons. By measuring the response threshold at each direction of the sound wave, we plotted the directional characteristics of each recorded auditory unit.

From the previous comparative physiological study on female mosquitoes (Lapshin and Vorontsov, 2013) we already knew that the auditory units of the JO are tuned to different frequencies. To correctly estimate the true auditory threshold of a given unit, its best frequency has to be measured first. Then, we used that frequency of stimulation sound to measure the directional properties of the same unit, i.e., the thresholds of its response to sound coming from different directions. Such a procedure, being repeated for a significant number of auditory units, gave us the overall directional characteristics of the JO. To adequately test for the hypothesised irregularities of these characteristics, both left and right JOs were studied.

On the next step, to describe the auditory space of a mosquito, which possesses two antennae, we needed to complement the individual auditory properties of each JO with the data on the relative orientation of the antennae. They are, however, not fixed, as insects, and mosquitoes in particular, can move their antennae in scape-pedicel joint by a set of muscles (Imms, 1939). To solve this task, we photographed flying mosquitoes and measured the orientations of their antennae in two planes and estimated the variation of these parameters. The combination of these measurements with the results on the directionality of individual auditory units implied a three-dimensional representation of the data, which we achieved by designing a graphical computer model to present the results in a simplified and easy-to-grasp form.

As the final goals of the study, we expected to understand (i) how a female mosquito could perform a binaural localization of a sound source based on the antennal directionality, and (ii) are there any regions of the auditory space of the mosquito to which it pays special attention in terms of sensitivity or resolution. We believe that the answer to the second question will lead to the design of more adequate behavioural experiments on female mosquitoes.

## 2. Methods

The method of measuring the directional properties of the JO sensory units was reported in detail in our study previous study on male mosquitoes (Lapshin and Vorontsov, 2019), here we describe it briefly and focus on the details specific to this study.

### 2.1. Animals

Females of *Culex pipiens pipiens* L. were captured in the wild in Moscow region of the Russian Federation. Experiments were performed at the Kropotovo biological station (54° 51’ 2” N; 38° 20’ 58” E) in August-September 2016-2021.

### 2.2. Behavioral experiments

Imaging of freshly captured mosquitoes was performed in the field, as we noticed that mosquitoes kept in laboratory for one or two hours flew continuously for significantly shorter periods. We placed individual mosquito into a circular plastic container, diameter 85 mm, depth 26 mm. Its flat walls were left transparent, while the curved side walls were covered with non-transparent film. The container was positioned either vertically (viewed from a side) or horizontally (viewed from above). In the former position, it was lit either by sunlight or by a home-made white LED array, intensity ca. 2000 lm, both through a light-scattering paper to provide a uniform bright background for imaging of a mosquito in a brightfield. The other flat side of the container was attached to a tube, which in turn was connected to a photo camera with a macro lens (Olympus OM-D E-M10 II, 60mm f/2.8). The pitch and roll of the camera were minimized relative to the horizon by means of a tripod. When viewed from above, the container was lit by scattered sunlight, and the same camera mounted on a tripod was used for imaging.

Mosquitoes (n=22) were serially photographed during the periods of flight, either spontaneous or caused by a startle from a shadow. From the images acquired in lateral view, we selected those where a mosquito was viewed strictly laterally (the second antenna was invisible) and was more than 1 cm from each wall. In the selected set of images (3 per each mosquito), the following angles were manually measured using the FIJI software package (Schindelin et al., 2012): antenna to horizon; head to horizon; abdomen to horizon, between the two antennae (in the view from above, projection to the horizontal plane).

From the measurements in lateral view we knew that the flying mosquito holds antennae at an angle to the horizon. When viewed from above, this angle does not allow to directly measure the inter-antennal angle. One way to overcome this issue would be to photograph a mosquito in two projections simultaneously. However, as we required only the average value of the inter-antennal angle, we corrected it using the average value of antenna-to-horizon angle according to the equation, derived from the geometry of the system:

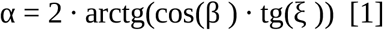

where α is a true inter-antennal angle, ξ is a half of the measured inter-antennal angle, projected to the horizontal plane, and β is an angle between the antennal plane (a plane formed by the two antennae, Fig.1A, dashed blue line) and the horizon.

**Figure 1.**
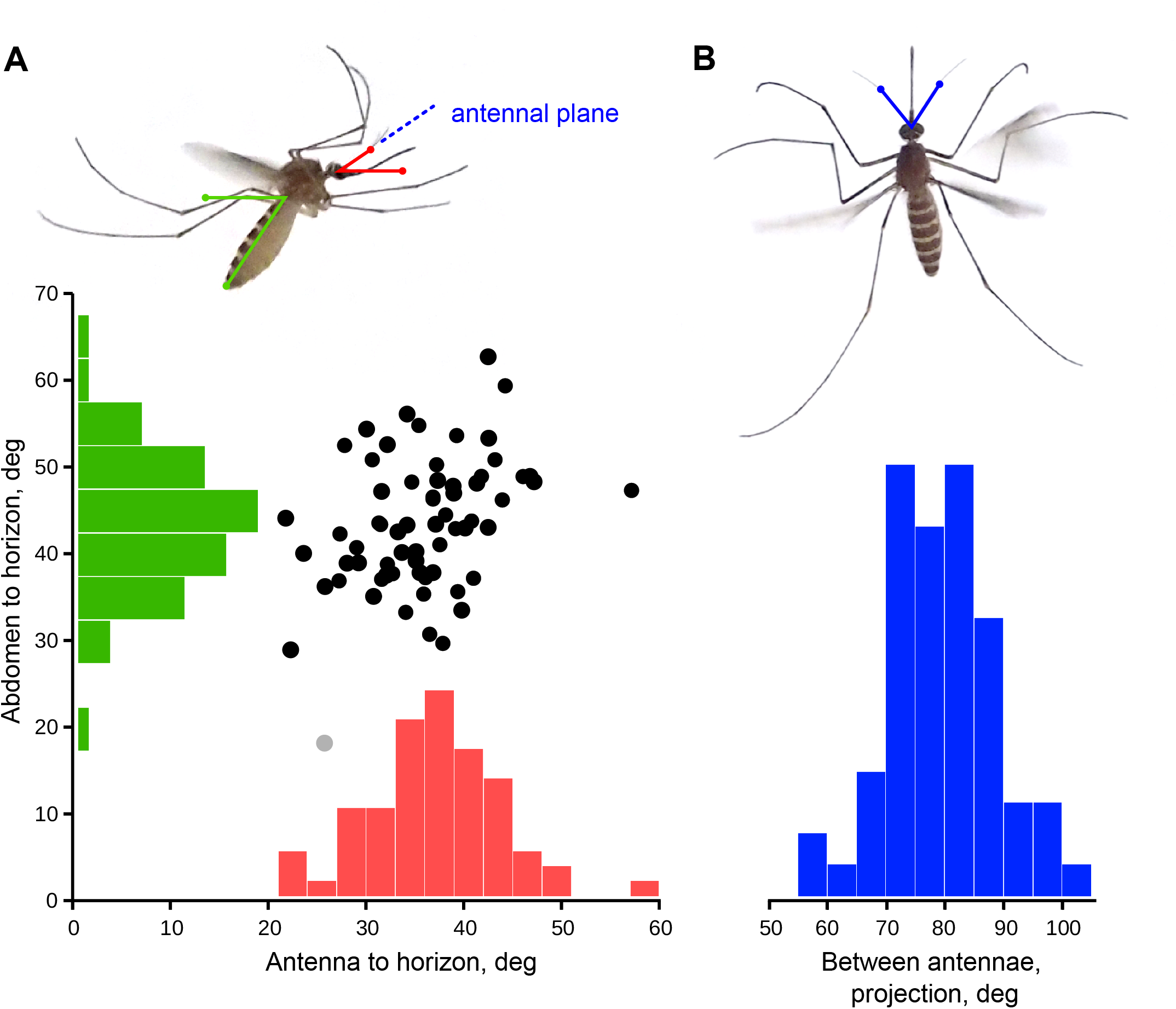
Orientation of antennae in a flying mosquito. The measurements were taken from multiple still images of flying mosquitoes. There was no clear dependence between the angles of antenna-to-horizon (red) from the abdomen-to-horizon (green), suggesting that the orientation of antenna to the world rather than to the body matters to mosquito. The true inter-antennal angle was calculated from the measurements in horizontal projection (blue) according to the equation [1] (see the Methods).

### 2.3. Microelectrode recordings

Experiments were conducted in laboratory conditions with air temperature 17–24°C. Focal extracellular recordings from the axons of the antennal nerve were made with glass microelectrodes (1B100F–4, WPI Inc.) filled with 0.15 M sodium chloride and inserted at the scape-pedicel joint. After the penetration of the cuticle electrodes had a resistance of 10–40 MΩ.

While penetrating the antennal nerve by the electrode, the preparation was continuously stimulated with tonal pulses (filling frequency 100–130 Hz, amplitude 60 dB SVPL, duration 80 ms, period 600 ms and dorso-ventral direction of acoustic vector (0°). During this searching procedure the groups of the JO neurons situated orthogonal to the antenna oscillation could be overlooked, so the vector of acoustic wave was periodically changed by 90°. We considered the recording site acceptable when the amplitude of response increased above 0.5 mV (peak to peak, Fig.2A). After that, the stimulation setup was switched into the feedback mode (see below).

**Figure 2.**
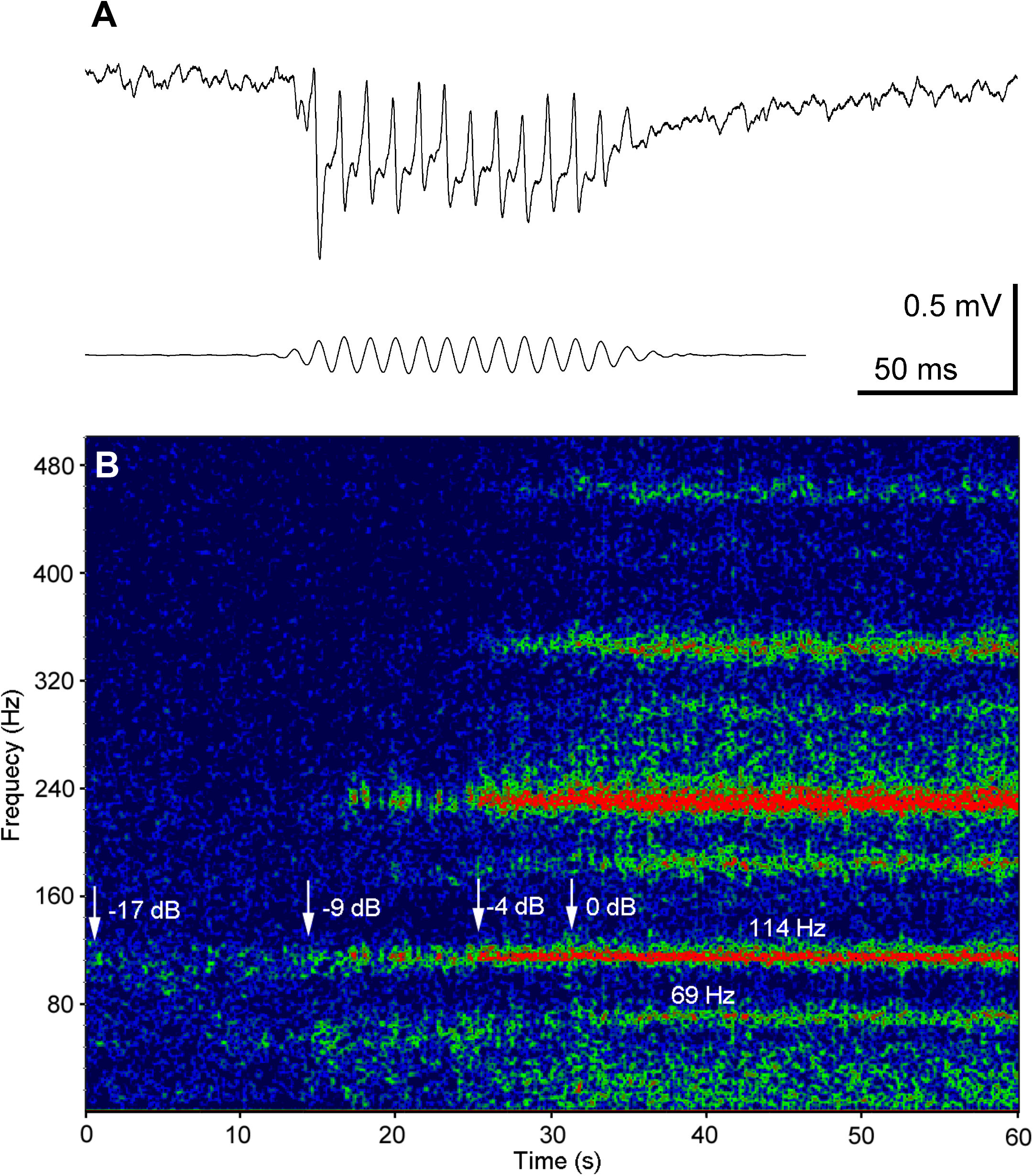
Responses recorded from the axons of auditory PSNs in the antennal nerve to different kinds of acoustic stimulation. A. Typical response to sinusoidal stimulation: glass microelectrode recording. Voltage scale is given for neuronal response. Lower trace: stimulation signal. (120 Hz, 58 dB SPVL), recorded by the microphone. B. Sonogram (frequency spectrum of signal against time) of response to a positive feedback stimulation. Colour represents the relative amplitude of response, from blue to red. The level of feedback was increased starting at −17 dB relative to the threshold (0 dB) established for this recording. The traces of the feedback effects appeared at −9 dB (narrow-band selective increase of noise). Note the two traces of simultaneous excitation at different frequencies (69 and 114 Hz).

### 2.4. Measurements of directionality

We measured the individual directional characteristics of the sensory units of the JO. In 162 experiments, of which 24 from the right JO, totally 290 (50 from the right JO) individual characteristics were obtained. For each recording site, two kinds of acoustic stimulation were used: positive feedback and sinusoidal. The former was used to obtain the individual tuning frequencies and unipolar directional properties of auditory units (hereafter ‘polar patterns’), while the latter allowed to measure the absolute auditory thresholds of a given unit at its tuning frequency and to plot the bipolar direction-threshold characteristics (hereafter ‘directional characteristics’).

|  | Left JO | Left JO,<br>low-frequency<br>search (70 Hz) | Right JO | Total |
| --- | --- | --- | --- | --- |
| number of mosquitoes/<br>experiments | 112 | 26 | 24 | 162 |
| number of individual polar<br>patterns (number of AFs) | 240 | – | 50 | 290 |
| number of individual directional<br>characteristics | 96 | 35 | 13 | 144 |

The essence of the positive feedback stimulation is a feedback loop established using the amplified in-phase response of a sensory unit as the signal to drive the stimulation loudspeaker. Applying such kind of stimulation to the sensory unit we expect it to ‘sing’ at the frequency which is close to its intrinsic tuning frequency – the effect hereafter called ‘autoexcitation’.

We used two orthogonally oriented stationary speakers to create a vector superposition of acoustic waves at the point of mosquito antenna. This approach enabled the rotation of the acoustic vector around the flagellum of the mosquito’s antenna. The mosquito was positioned at the crossing of the axes of two speakers in such a way that the antenna′s flagellum was perpendicular to the directions of sound waves originating from each of the two speakers. In the angular coordinates used throughout this study, φis the angle between the dorso-ventral line passing through the mosquito′s head and the vector of vibration velocity of air particles. An increase in φcorresponds to counter-clockwise rotation of the velocity vector, with the insect’s head viewed frontally. Accordingly, when viewed from the mosquito′s head along the antenna (Fig.3, inset), the clockwise rotation corresponds to the increase in φ.

**Figure 3.**
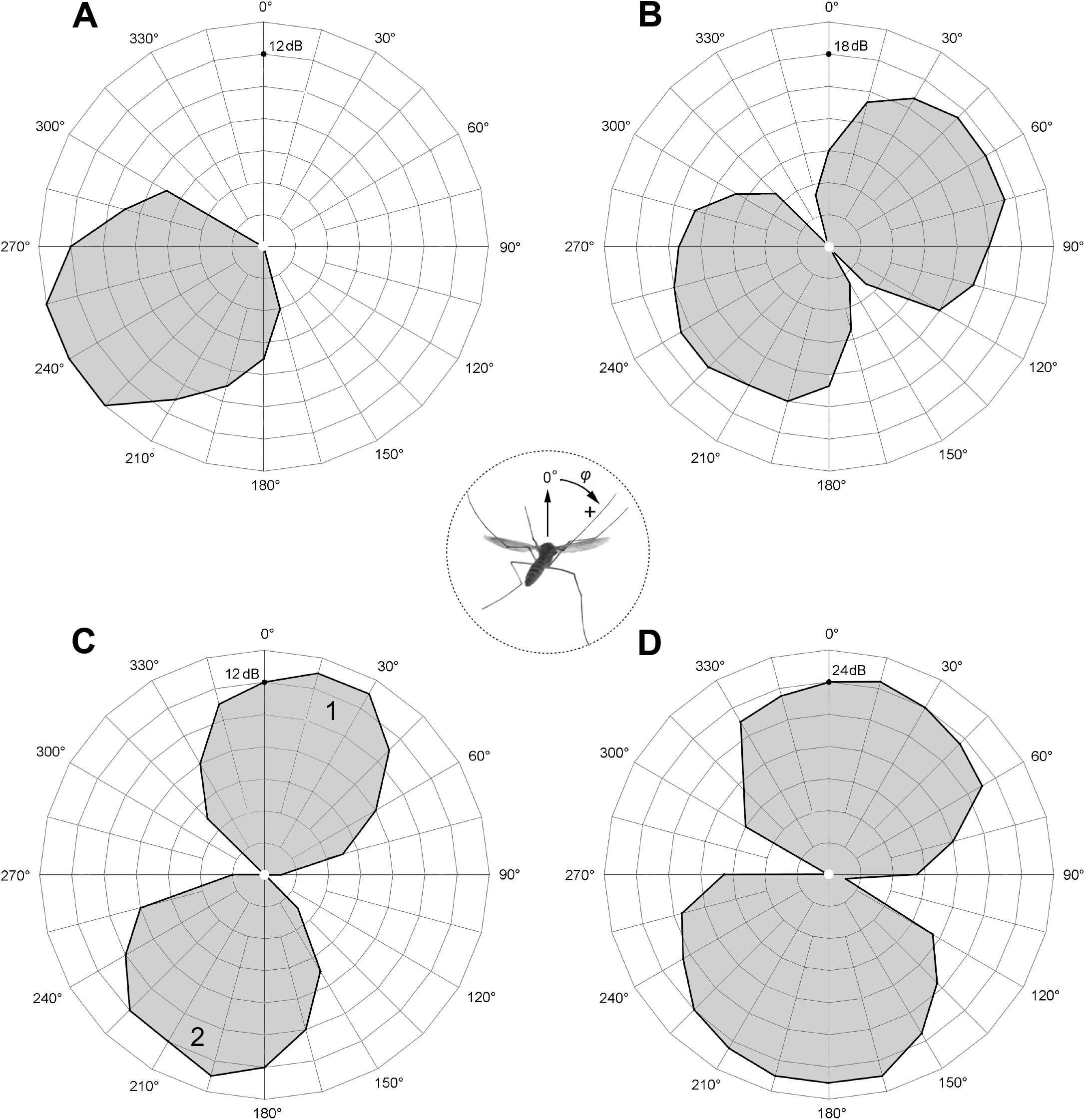
Examples of directional diagrams of the JO auditory units. A. Polar pattern of a single responding unit (AF 112 Hz). B. Directional diagram of the same unit measured by sinusoidal stimulation; threshold of response 32 dB SPVL at 112 Hz. C. Polar patterns of two units recorded at the same site and responding in anti-phase; the tuning frequencies are 104 Hz (unit 1) and 77 Hz (unit 2); D. Directional diagram of the same pair of units as in C, measured as response to 100 Hz sinusoidal stimulation; absolute threshold 38 dB SPVL. Image of mosquito viewed posteriorly shows the positive direction of angular axis used in all diagrams for both left and right JOs.

The stereotyped way of positioning the mosquito relative to the experimental setup, especially to the recording electrode, could lead to selective recording only from a certain part of the antennal nerve, as the final search for the recording site was performed blindly with the electrode tip hidden below the cuticle. To minimize the probability of such bias, the mosquito was occasionally positioned ventral side up (180° rotation, n=15) or rotated by ±45° (n=16) or ±90° (n=37) around the longitudinal axis of its body. In each case, the recording electrode was inserted through a different part of the scape-pedicel joint. The directional data measured in all such experiments were corrected accordingly. To make an additional control of the above-mentioned experimental bias, in a separate series of experiments (n=24) we performed recordings from the right JO instead of the left one, keeping similar all other properties of the experimental setup. To avoid confusion due to the possible symmetrical nature of the two antennae, unless specifically stated, the presented data include measurements from only the left JO, while the data from the right JO serve only as a control.

### 2.5. Search for low-frequency units

Based on preliminary results, we realized that the true responses of low-frequency sensory units are often difficult to distinguish from combination harmonics that appear when two or more high-frequency units respond simultaneously at different frequencies (example in Fig.2B, trace at 69 Hz). This effect could lead to the under-estimation of true low-frequency-tuned units. Also, the method of positive feedback stimulation was found to be rather ineffective at the lower frequencies, most probably, due to the high competition from the more sensitive units tuned to higher frequencies, which often captured the autoexcitation frequency. Accordingly, we designed an additional series of experiments to compensate for these biases. In these experiments, the searching frequency was set to 70 Hz, and during the choice of the recording site we gave preference to the units that responded maximally at that or at lower frequencies. Moreover, the initial (searching) vector of the acoustic wave was set to φ= −45°, that is, to search for units in II and IV quadrants, where, according to the preliminary analysis of the data, we expected to find less responding units.

### 2.6. Measurements of thresholds

In the mode of feedback stimulation, we could only measure the relative threshold of auto-excitation of the entire system, including the mosquito and the stimulation setup. Such a relative threshold was defined as the signal level, which required one more incremental step (+1 dB) at the attenuator output for the system to enter the state of sustained autoexcitation (Fig.2B).

After finding the best direction using the positive feedback stimulation we measured a frequency-threshold curve in the range 40–300 Hz using sinusoidal stimulation with a 5 Hz step below 170 Hz and a 10 Hz step otherwise. The criterion of the response threshold was set at 2 dB of sustained excess of response amplitude above the average noise level in a given recording. At each combination of stimulation parameters the threshold was measured consecutively at least twice. Below 40 Hz, the measurements were limited by substantial decrease of efficiency of the speakers. The level of stimulation was limited by 80 dB SPVL to avoid overloading of mosquito’s auditory system by too loud stimulation.

Similarly, after finding the best frequency of a given unit through the positive feedback stimulation, we measured a direction-threshold plot (auditory threshold as a function of orientation of the acoustic wave vector relative to the axis of the antenna) with a 15° step, using sinusoidal stimulation.

### 2.7. Data processing

#### Directional plots

In an array of threshold data obtained from a single recording site, we determined the maximum threshold value (Th _max_). Based on it, a set of derived values describing each unit’s directional characteristic or polar pattern was estimated by the formula A_i_ = Th _max_ – Th _i_. In the plots based on these data, the sectors of the highest sensitivity corresponded to the lowest recorded thresholds, and the central zero point corresponded to Th _max_. The angles at which no response at the best frequency was observed were given the value A_i_ = 0.

The angular sensitivity range of a unit was determined at −6 dB of its maximum sensitivity (in case of directional characteristics the values from the two independently measured symmetrical plots were averaged). The best direction of a given unit was determined as the bisector of this range.

#### Individual tuning frequencies

Using the positive feedback stimulation, we determined the individual autoexcitation frequencies (AFs). The value of the excitation frequency was inevitably affected by the total phase shift in the feedback loop, arising primarily from the latency of the sensory cell response. As the phase shift was not constant along the frequency range, for each frequency measurement, a compensatory phase shift was introduced into the loop. The value of that phase shift was selected to provide the maximal amplitude and stability of autoexcitation at the same level of feedback.

#### Correction for temperature

As the experiments were performed in a range of ambient temperatures, all frequency data were normalised to 20°C according to the previously established formula: F_20_=F_t_ (1+k (20-t)), where t is a temperature for a given experiment and k=0.02 is a relative shift of tuning frequency with a change of frequency by 1°C (Lapshin and Vorontsov, 2021).

## 3. Results

### 3.1. Position of the antennae in flying mosquitoes

The angle at which a flying mosquito holds its antennae relative to the horizon was 35.5° on average (n=66), with no clear dependence on the angular position of the body, measured along the abdomen (Fig.1A). The inter-antennal angle, directly measured as the projection to the horizontal plane, was 76.8° (n=63, Fig.1B), which corresponds to 73.9° in the plane of the antennae, according to the equation [1, Methods].

### 3.2. Individual directional characteristics of sensory units

A typical threshold directional characteristic of an individual sensory unit measured with the sinusoidal pulse stimulation was symmetrically bi-directional, showing a classic figure-eight pattern (Fig.3BD). The width of each petal was 121° (σ=18.6°, n=96).

Acoustic stimulation in the positive feedback mode allowed, as in previous studies (Lapshin, Vorontsov, 2019), to discriminate several different responses at each recording site. First, the feedback stimulation combined with directionality test revealed uni-directional polar patterns (Fig.3A). With slight deviations (±8°) they overlapped with one of the petals of the bi-directional characteristics measured at the same site with sinusoidal stimulation (Fig.3B). These results were fully expected, as one and the same sensory unit should demonstrate exactly such kind of responses: the phase of the stimulation matters in the positive feedback stimulation, but not in the sinusoidal mode, where a unit cannot discriminate the two opposite directions. This should not be confused with the two different units, each possessing individual frequency tuning, responding at the same recording site (see below).

In 54 recordings (23% of the total, n=240) there existed only a single uni-lateral response to a feedback stimulation, not complemented with other responses at different phases (Fig.3A).

In most recordings (n=162, or 67%) there appeared another response with a different AF (Fig.3C). Its polar pattern was oriented oppositely (180±10°) to the first one. In such recording sites with an anti-phase pair of units the sinusoidal stimulation revealed similar thresholds of response (ΔTh≤3 dB) to both the AFs determined by feedback stimulation or to the intermediate frequency (Fig.3D).

In rare cases, there was simultaneous auto-excitation at two different frequencies within a single polar pattern and at a third frequency in the opposite direction (n=6) or without any response in the opposite angular range (n=3).

### 3.3. Radial distribution of sensory units in the JO

Fig.4A shows the distribution of sensory units from the left JO depending on their best directions. The same data are plotted in the polar coordinates centered on the axis of the left antenna in Fig.4C. The units are distributed unevenly: the number of units oriented in the first and third quadrants is 3.2 times higher compared to the second and forth quadrants (181/56, χ^2^=33, p<0.001). The same bias persisted in the subset of the data when the mosquito was initially rotated by 90° relative to the stimulation system and the recording electrode holder (Fig.4A, black-filled part of the distribution, angular data corrected respectively). The control series of measurements made from the right JO revealed most units oriented in the second and forth quadrants (31/9, χ^2^=6.05, p<0.025, Fig.4B), i.e. mirroring the distribution obtained from the left JO. Results of both controls speak in favour of true asymmetry in the distribution of sensory units in the JO, characteristic of *Culex pipiens* mosquitoes.

**Figure 4.**
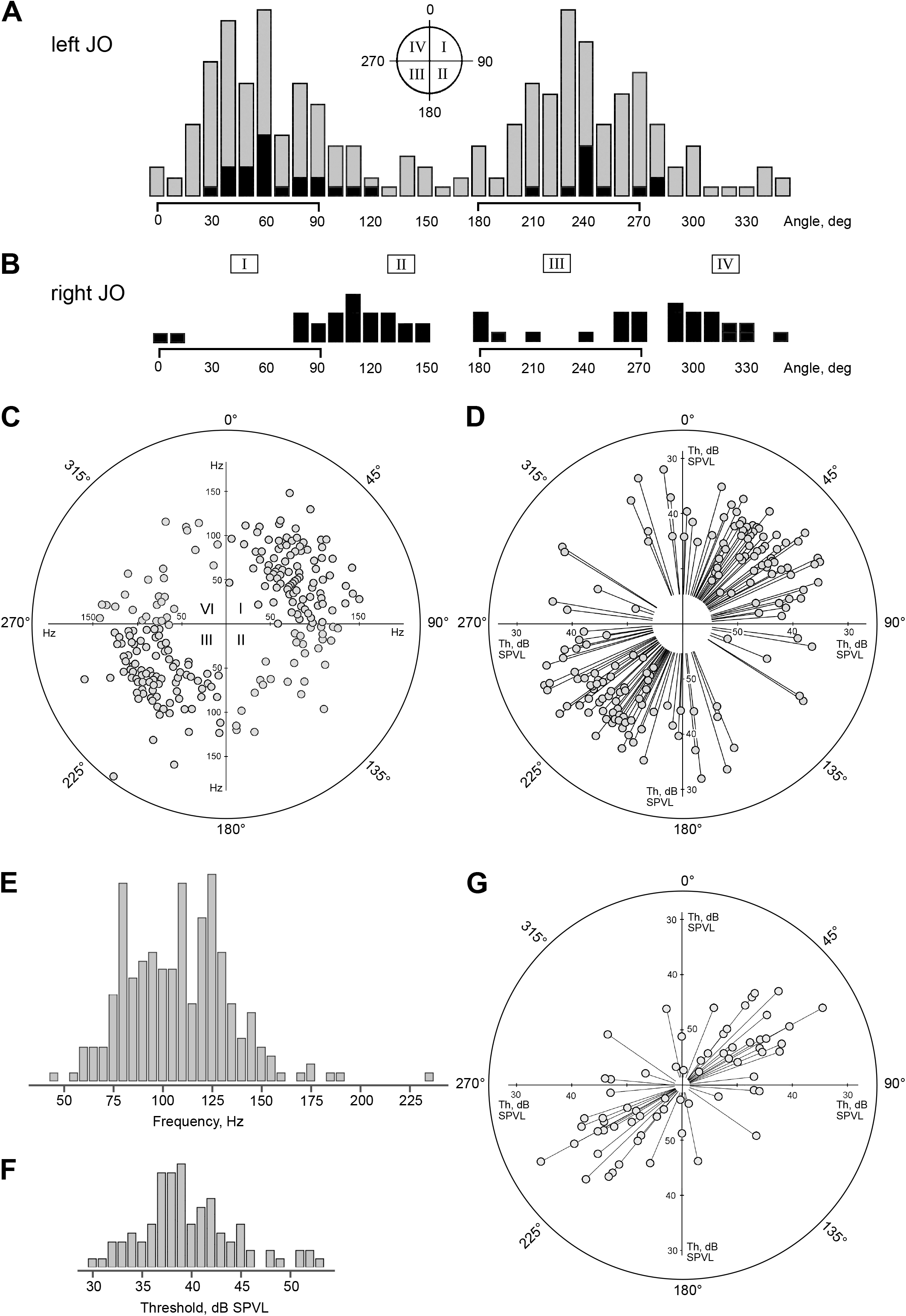
Distributions of auditory units around the axis of the JO. A.Left JO, angular orientation similar to Fig. 3. Gray bars show overall distribution, black bars represent the data from experiments where a mosquito was positioned ±90° around its antero-dorsal axis (the angular data were corrected accordingly). Note that, regardless of mosquito orientation and the position of inserted electrode relative to the head and the JO, the majority of sensory units belong to the I and III quadrants (0–90 and 180–270 degrees). B. Similar data obtained from the right JO. Note that the distribution mirrors that for the left JO, i.e. right auditory units are oriented in the II and IV quadrants. C. Polar distribution of individual frequencies (AFs), left JO. In 11 cases the points fully overlapped each other. D. Polar distribution of individual thresholds, left JO. Lower thresholds (higher sensitivity) are further from the center. E. Distribution of individual tuning frequencies (AFs) measured by a positive feedback. F. Distribution of auditory thresholds of sensory units, each measured at the best frequency for a given unit. G. Polar distribution of individual thresholds of low-frequency-tuned units (below 80 Hz), left JO. Note that these units are oriented similarly to the higher-frequency ones (D), although the initial orientation of the searching stimulation signal favoured the finding of units oriented in the two alternate quadrants (II and IV).

In the distributions of best directions of units in both left and right JO’s (Fig.4AB) there are two additional sub-peaks at 80–90° and 260–270°. In other words, 17 % (n=42) of units were oriented in the antennal plane and their proportion was significantly higher than that of the units oriented perpendicular to the antennal plane (N_h_=42, N_v_=15; χ^2^=6.4, p<0.025).

Sinusoidal stimulation allowed to measure the absolute thresholds of units, which are plotted in polar coordinates in Fig.4D. Due to the symmetrical nature of the directional characteristics, each data point on the plot has its pair situated symmetrically relative to the center of the coordinates.

The individual thresholds of recorded units varied from 30 to 53 dB SPVL, with 39.8 dB SPVL on average (σ=4.8 dB, N=96), their distribution is shown Fig.4F. Most sensitive units with thresholds of 36 dB SPVL or lower were found around 53° (Fig.4D) with tuning frequencies in the range 65–135 Hz.

The data from a separate experimental series (70 Hz, −45° searching stimulation, which gave preference to lower frequency units oriented in second and fourth quadrants) also demonstrates that most recorded units was oriented in the first and third quadrants (27/6, χ^2^ = 6.68, p<0.01, n=25, Fig.4G), i.e., similarly to the main array of data for the left JO (Fig.4D).

### 3.4. Audiograms

Most of the audiograms measured from either the left or right JO can be divided into three groups by the position of the main minimum (the range of highest sensitivity): i) main minimum at 90–100 Hz or higher. This group represents most cases (n=127, 78%, Suppl. Fig.1A, black curve); ii) main minimum at 50–60 Hz (n=13, 8%, Suppl. Fig.1A, green curve); iii) no obvious minimum, with a slope towards the lower frequencies (n=22, 14%, Suppl. Fig.1A, blue curve). The measurements of thresholds at even lower frequencies were limited by the stimulation equipment.

In most cases, it was difficult to isolate individual responses of a unit from a compound response to measure the Q-factor of a unit. However, we managed to do it in several recordings of higher quality: examples of audiograms measured in such experiments are shown in Suppl. Fig.1B together with Q-factor values estimated at +6 dB from the minimal thresholds.

The distribution of auto-excitation frequencies (Fig.4E) is neither normal nor log-normal (Pearson test, p>0.05). It contains two local components that differ from the theoretical curve: minimum at 10^**1.94**^=87 Hz (χ^2^=4.03; p<0.05) and maximum at 10^**2.1**^=126 Hz (χ^2^=11.24; p<0.001).

### 3.5. Ratios between individual frequencies in pairs and triplets of units

The histogram of the ratio between individual frequencies in pairs of anti-phase units F2/F1 (F2>F1) has a distinctly regular structure (Suppl. Fig.2A, n=81). The distribution is asymmetric as it is limited below 1. Taking the logarithm of the data removes this restriction (Suppl. Fig.2B), but still the resulting data did not pass the Pearson test for normality (χ^**2**^ (0.05; 60) = 79 < Σ χ^**2**^=174). The sum of main components Σ χ^**2**^ corresponds to the four major peaks of the original distribution: F_**2**_/F_**1**_=10^**0.096**^=1.25 (χ^**2**^=8.77; p<0.01), F_**2**_/F_**1**_=10^**0.125**^=1.33 (χ^2^=30; p<0.001), F_**2**_/F_**1**_=10 ^**0.175**^=1.5 (χ^2^=31; p<0.001), F_**2**_/F_**1**_=10^**0.225**^=1.67 (χ^**2**^=28; p<0.001). The values of the peaks are either equal or close to the integer fractions 5/4, 4/3, 2/3, and 5/3. We had not enough data for triplets of units (n=6) to perform similar analysis, however, the ratios between individual frequencies demonstrated the same pattern.

## 4. Discussion

### 4.1. Directionality of sensory units in three dimensions

Positive feedback stimulation enables the separation of the activity of two antiphase units in paired systems (for detailed discussion, see Lapshin and Vorontsov, 2019). Under sinusoidal stimulation, both units possess similar sensitivity to acoustic signals coming from the opposite directions (Fig.3BD). In other words, sensory units with polar patterns oriented in the 1st quadrant will also receive signals from the directions of the 3rd quadrant, and vice versa. The same is true for the units oriented in the 2nd and 4th quadrants.

The resulting three-dimensional audiogram of a single JO sensory unit is combined from the mechanical directional diagram of the antenna and the characteristics of a unit in a plane perpendicular to the antenna. The diagram of the antenna in the first approximation has the shape of a torus with minima on the apex and base and a maximum in the plane perpendicular to the flagellum (Fig.5A). An averaged two-dimensional directional characteristics of a unit consists of two symmetrical petals of more or less circular shape (Fig.3BD). Then, the simplified three-dimensional characteristics of a single sensory unit should be a system of two contacting spheroids with their common axis oriented at an angle (*φ)* to the transverse axis of the JO.

**Figure 5.**
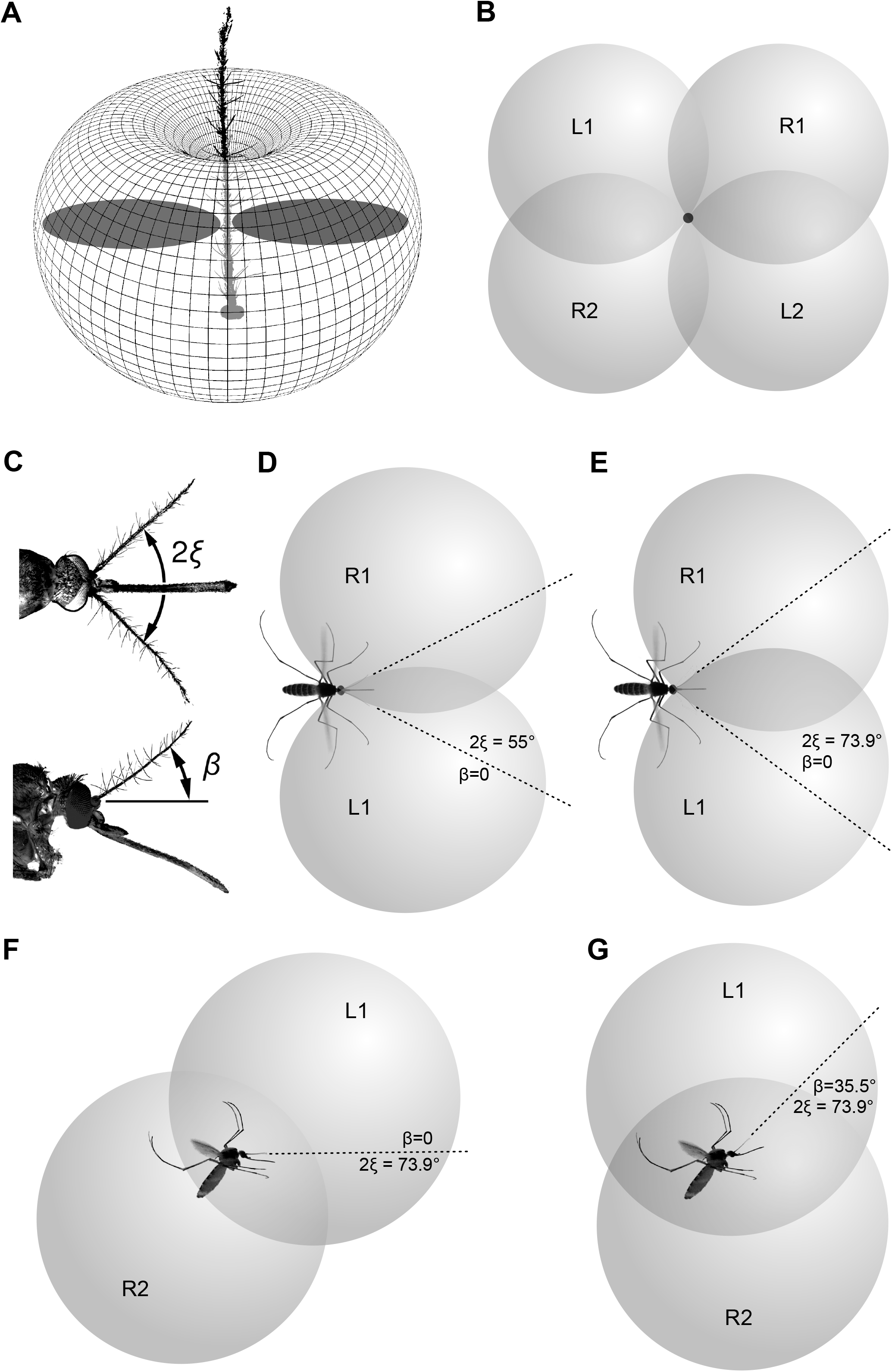
Schematic illustration of three-dimensional directional characteristics of the two symmetrical groups of sensory units from the left and right JOs, oriented at φ=53°. A. Theoretical directional diagram of the mosquito antenna. It has the shape of a torus with minima on the apex and the base of antenna and a maximum in the plane perpendicular to the antenna. Flat directional diagrams of an individual unit, as measured in this study, are shown in dark-grey. B. Diagrams of left (L1, L2) and right (R1, R2) sensory units when antennae are parallel and oriented towards the viewer (this is non-existent simplification provided for step-by-step explanation) C. Actual position of antennae in flying mosquito (see also Fig. 1). Antennae are deflected from the parralel in frontal (angle 2*ξ*) and sagittal (angle *β*) planes. D, E. Intersection of left and right diagrams when antennae are spread apart (but not raised, *β*=0), at different values of an angle between antennae (2*ξ*). dorsal view. Parts of diagrams L2 and R2 not shown for simplicity. F. The same, 2*ξ =* 73.9*°, β*=0 lateral view. G. Resulting position of directional diagrams with orientation of antennae as shown in C (2*ξ* = 73.9*°, β*=35.5°), lateral view. Dark-grey intersection areas in B, D and E show equisignal zones where the ratio between the responses of the left and the right units does not depend on the absolute level of sound.

Binaural hearing would require the comparison of responses coming from the left and the right antennae. In terms of their 3D directional diagrams, the transition from a single sensory unit to a set of units belonging to different antennae may be difficult to imagine. Below we will explain this transit in sequential steps. In the Supplementary data we provide a video file and a parametric computer model that graphically illustrates the steps described below.

For the sake of initial simplicity, let us temporarily assume that the two antennae are held horizontally and in parallel to each other. Then, characteristics of the two symmetrically oriented units, belonging to the sectors that are populated with the most sensitive ones, φ=53° on average (Results, 3.3, φ= −53° for the right JO), will overlap like in Fig.5B, mosquito viewed from the front, characteristics L1+L2 and R1+R2 belong to the units of left and right antennae, respectively.

The antennae of female mosquitoes in their average in-flight position are spread apart and form an angle, 2ξ (Fig.5C). Given such geometry, the parts of directional characteristics L1 and R1 shift forward and medially, while L2 and R2 shift backward and medially (Fig.5DF). With such a shift, the overlap between the left and the right characteristics increases proportionally to the angle 2ξ (Fig.5DE).

The upward inclination of the antennal plane (angle β) relative to the horizon (Fig.5C) can compensate for the backward-forward displacement of characteristics at the previous step: their dorsal parts (L1 and R1) shift backward, while the ventral ones (L2 and R2) shift forward (Fig.5G). Geometrically, the condition of full compensation, when the dorsal auditory space is located strictly above the ventral one, is described by the following equation:

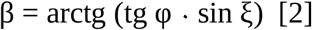

Given the measured inter-antennal angle in flying mosquitoes, 2ξ=73.9°, for the most sensitive units oriented at 53° the angle β will be 38.6°. This is very close to the measured value of β, 35.5°.

It may seem counter-intuitive, but the spreading of the antennae apart allows a mosquito to significantly increase, rather than decrease, the area of overlap between the diagrams of left and right groups of most sensitive auditory units, which may enhance binaural hearing (Fig.5DE, see also supplementary video). The upward inclination of the antennae may compensate for the skewness of the overall auditory space (Fig.5FG).

Since the best direction (φ) has been varying among the JO auditory units, there is no position of antennae that would be optimal for all units. Their angular distribution (Fig.4CDG) together with the most frequently observed orientation of antennae (Fig.1) suggest that female mosquitoes pay special attention to the sounds coming from above and below. However, as the position of the antennae may vary in different behavioural states, a mosquito can constantly adjust the parameters of its directional hearing depending on the current sensory task.

### 4.2. Equisignal zones

The most remarkable feature of the overlap between the left and the right directional characteristics is that it forms the so-called equisignal zones (shown in dark-grey in Fig.5DE). Within these zones, the angular position of a sound source relative to a mosquito can be estimated from the ratio between the amplitudes of response coming from the left (A_left_) and the right (A_right_) units:

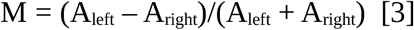

The functional advantage of the parameter M is that it does not depend on the absolute level of sound. However, such a simplified scheme does not allow to unambiguously determine the direction to the sound source: the sounds coming from each of the four equisignal zones (Fig.5B) can result in similar ratios of responses in A_left_ and A_right_. For a more precise localization of the sound source, a mosquito should analyse a more complex set of signals produced by auditory units oriented at different angles within each JO. In addition, the acoustic directionality of a flying mosquito must be aided by its active maneuvers. Remarkably, the overlap of directional characteristics of the left and right antennae, at least those of the units oriented at φ=53°, becomes higher when antennae are spread apart than when they are brought together (Fig.5DE).

As already stated, due to the symmetry of a unit’s directional diagram relative to the antenna flagellum (Fig.3B), there exists an uncertainty in the localization of the sound source in the two opposite directions. However, earlier Belton (1974) pointed out that the phase of oscillations in both antennae may be significant for directionality. A point source of sound at an angle of greater than 40° from the antennal plane would deflect both antennae in phase, while a sound source directly in front of the antennae would deflect the two flagella out of phase. It is quite possible that the brain of a mosquito performs a combined analysis of the amplitude and phase ratios of the signals coming from the right and left JOs to reduce uncertainty in estimating the direction to the sound source.

### 4.3. Functional asymmetry of the JO

All the evidence we have to date indicates that the auditory function of the mosquito JO is asymmetrical: although its morphology demonstrated radial symmetry (Boo and Richards, 1975; Hart et al., 2011), most of the units responding to sound were found in two of four sectors (sectors I and III of the left JO, Fig.4ACDG). Recording from the contralateral (right) JO revealed a similar but mirrored distribution (Fig.4B), contesting the hypothesis that the observed asymmetry was an artefact. At the same time, each JO possesses units oriented in two other sectors (II and IV) which, given the three-dimensional geometry of the two antennae, may duplicate the function of the contralateral ones.

The second largest group of units is oriented in the frontal plane (φ≈ 90° and φ≈ 270°, see sub-peaks in the distribution, Fig.4A). When the antennae are spread apart to the angle of 2ξ ≈ 74°, parts of the directional characteristics of these receptors are shifted forward, forming another wide equisignal zone directly in front of the mosquito. Inclination of the antennal plane (angle β) leads to rising of this zone into the upper hemisphere of the mosquito’s auditory space.

### 4.4. Frequency tuning

After finding that quadrants II and IV of the left JO are depleted in sound-responding units (Fig.4CD) we tested different hypotheses explaining such functional asymmetry in a morphologically symmetrical system. One of those was that these sectors of the JO are populated primarily by low-frequency units tuned below 80 Hz which we could skip due to inadequate frequency of the searching signal (100–130 Hz). Additional experimental series with low-frequency searching signal discarded this hypothesis: the radial distribution of low-frequency units did not show significant differences from a high-frequency one (Fig.4G). However, these experiments allowed us to study the low-frequency units which were indeed overlooked in the main series. According to the combined distribution of tuning frequencies (Fig.4E) and the number of recorded narrow-band units (Suppl. Fig.1B) the hearing of *Culex pipiens* female mosquitoes ranges from 40 to 260 Hz. Low-frequency units (40–80 Hz) were found 6 dB less sensitive on average, but the values of their minimal thresholds (ca. 30 dB SPVL) were similar to the units tuned above 80 Hz. Thus, the ability of female mosquitoes to hear low frequencies extends down to at least 40 Hz without a significant drop in sensitivity.

Earlier it was shown that the JO of Culex female mosquitoes possesses low-frequency sensory units that significantly decrease their thresholds (by 40 dB, or by 100 times, at 40 Hz) when affected by the signal simulating the flight vibration, i.e. when a second dorso-ventral acoustic stimulus was applied as a background (Lapshin, 2013). It is possible that female mosquitoes possess a specialized set of low-frequency-tuned auditory cells that are intended for in-flight reception of some low-frequency signals, such as sounds accompanying the movements of host animals.

What is the functional role of the sensory units oriented in the quadrants depleted in auditory neurons is still a question. Studies in *Drosophila* suggest that not all JO neurons in these insects are used for hearing: only about half the JO neurons responded to antennal vibrations and mediated hearing, while the other half detected the antennal deflections imposed by gravity and wind (Kamikouchi et al., 2009; Yorozu et al., 2009; Matsuo and Kamikouchi, 2013).

### 4.5. Interactions between the paired auditory units

The observed clearly discrete nature of the distribution of frequency ratios in paired systems of units (Suppl. Fig.2) indicates an electrical or mechanical interaction between the units in a pair. This phenomenon can be attempted to explain by the fact that auditory receptors of the JO in mosquitoes possess the mechanisms of endogenous activity, or generators, that help to amplify low-amplitude signals (Göpfert and Robert, 2001; Avitabile et al., 2010).

When two generators, tuned to different frequencies, become coupled, with high probability, this leads to a phenomenon called harmonic synchronization. Such kind of interaction manifests itself in the mutual adjustment of the frequencies of both generators in order to reach some integer ratio between the frequencies. This state corresponds to a local energetic minimum of the system and contributes to stability of harmonic synchronization (Yang et al., 2012). Thus, the observed mutual synchronization of auditory units in the mosquito auditory system may occur due to a mechanism which is common in nature and is not necessarily a sign of specific adaptation. Affected by some destabilising factors, a pair of coupled generators can switch to different modes of synchronization, for example, changing a frequency ratio from 3/2 to 4/3. In addition, the range of possible frequencies which a generator can be tuned to can have biophysical constraints, in that way additionally limiting the choice of a frequency ratio. Most probably, the combination of all these factors leads to the appearance of a set of integer frequency ratios in paired systems of mosquito auditory units: 5/4, 4/3, 3/2, and 5/3 (Suppl. Fig.2A). Morphologically, sensillae that include only a single receptor cell have not yet been described in the mosquito JO; all auditory sensillae contain at least two mechanoreceptor cells (Boo and Richards, 1975). Meanwhile, a significant probability of recording single units (23%) implies their wide representation in the JO. It can be assumed that the activity of such “single units” is in fact a synchronous response of two units tuned to the same frequency (F2 / F1 = 1) and belonging to a single sensilla.

The functional significance of paired auditory units may have different but complementary explanations. The system of two in-phase paired units may have improved signal to noise ratio, as the total noise of the system is summed up as a root-mean-square while the in-phase signals are linearly summed, thus increasing the signal-to-noise ratio of the paired system by 2/√2=√2, or 3 dB in terms of a threshold. The anti-phase pairs of units, which are the most abundant in the mosquito JO, may represent a system of selective frequency filters. In each aniphase pair, all oscillations beyond the frequency tuning ranges of the two units, including the low-frequency deflections caused by wind currents, get mutually subtracted.

## 5. Conclusions

Female mosquitoes possess a sophisticated system of directional hearing and, most probably, can adjust it by moving the antennae. The average in-flight position of antennae together with anisotropic distribution of auditory units in each JO suggests that the female mosquito pays primary attention to the areas strictly above and below itself. There is another area in front of and above the mosquito where directional characteristics of both antennae significantly overlap, providing binaural hearing. Spreading of the two antennae apart allows a mosquito to increase, rather than decrease, the overlap of the directional diagrams of most sensitive auditory units, thus enhancing binaural hearing. At the same time, each JO possesses units that duplicate the function of the contralateral one. Such redundancy increases the reliability of the entire system and potentially can provide acoustic orientation even if one antenna gets damaged.

## Supporting information

Supplementary computer 3D model

## 6. Acknowledgments

We are grateful to Ian Russell (University of Brighton, UK) and Patricio Simões (University of Sussex, UK) for valuable discussion. The field facilities for this study, Kropotovo biological station, was provided by the Koltzov Institute of Developmental Biology RAS.

## 9. Supplementary files

**Supplementary Figure 1.**
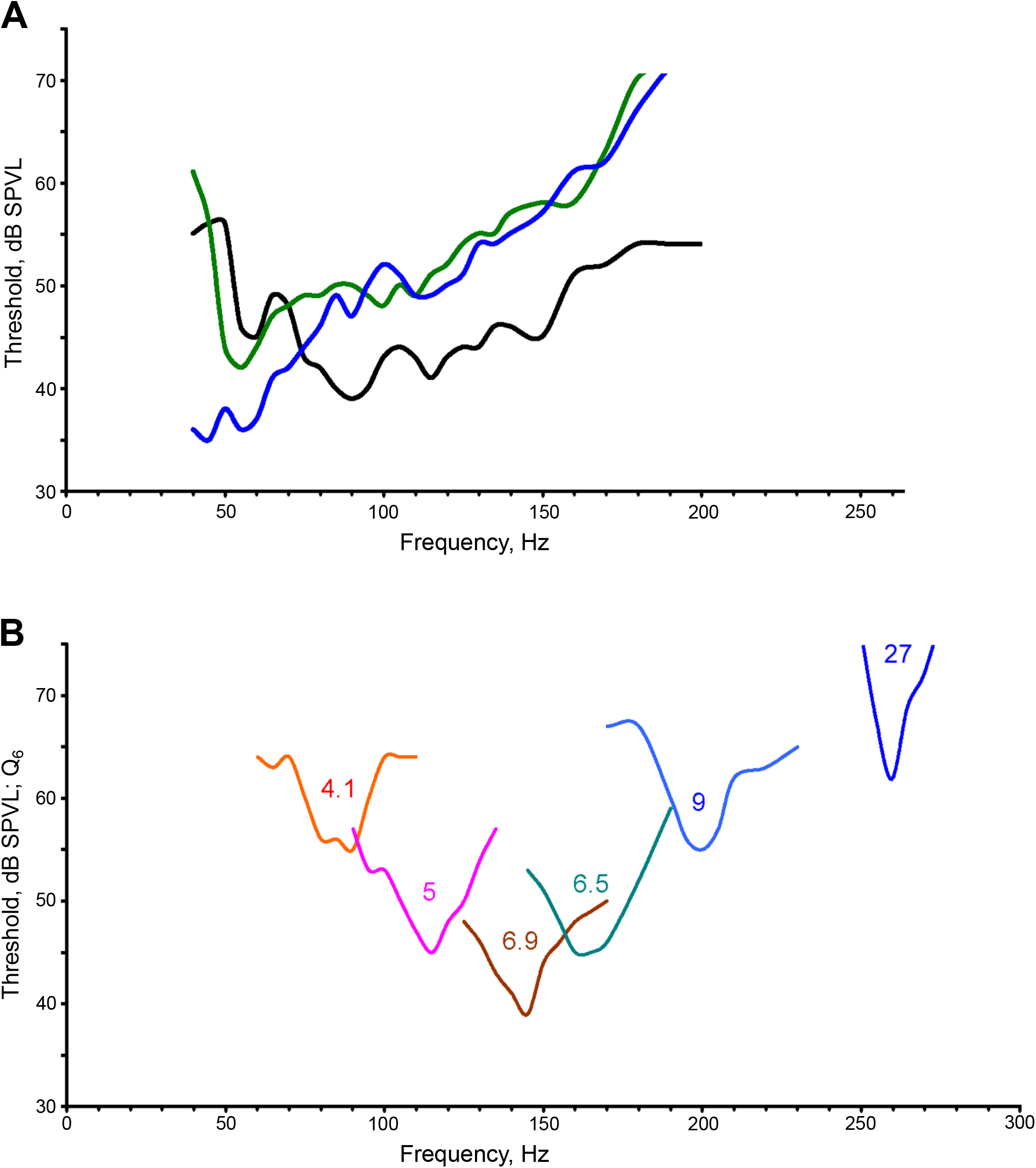
Examples of frequency-threshold curves (audiograms) of the JO auditory units. A. Black curve represents the majority of recorded units, with best thresholds at 90–100 Hz or higher. Green curve, best thresholds at 50–60 Hz. Blue curve, slope towards the lower frequencies, (the tested frequency range was limited by the stimulation equipment). B. Individual curves of narrow-band units measured with a 5 Hz step. Numbers show quality values (Q_6_, measured at +6 dB from the best threshold). Note that such narrow-band units cover almost the whole frequency range of female mosquitoes (40 to 260 Hz) and that their thresholds are mostly not among the lowest ones (compare to Fig.4F).

**Supplementary Figure 2.**
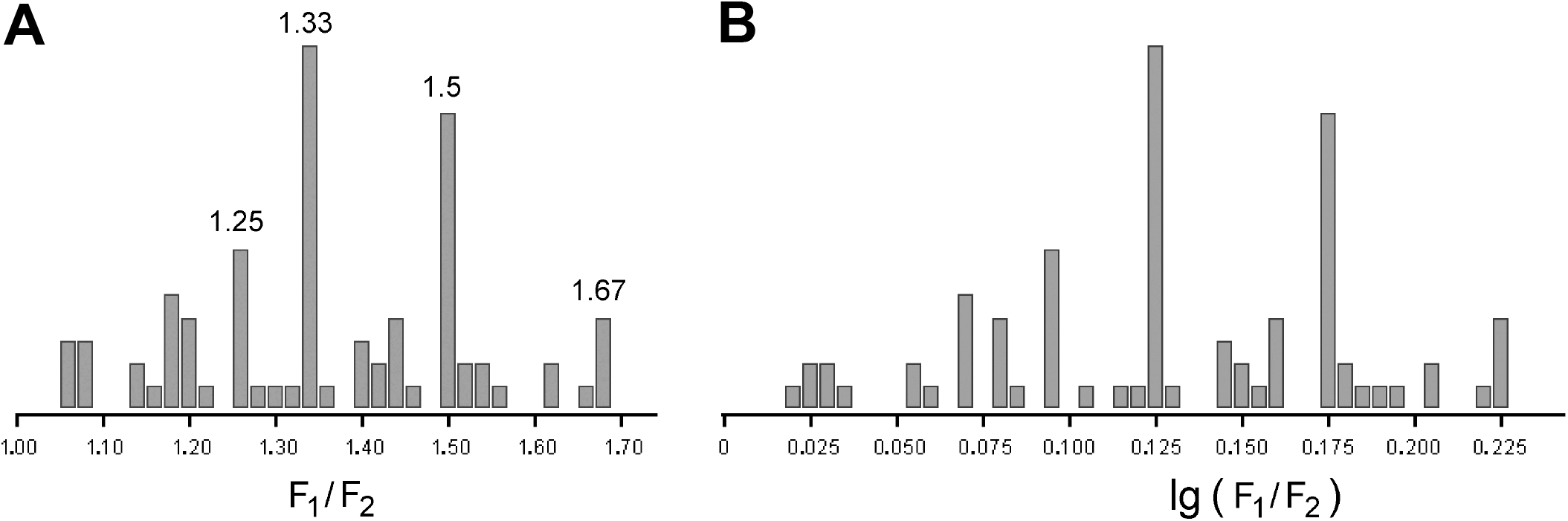
Ratios of tuning frequencies in paired auditory units. A. Numbers show the values of the most prominent peaks, which correspond to the integer ratios (3/2, 4/3, 5/3, 5/4). B. The same data as in A, logarithmic scale

**Supplementary video**. Screencast of a supplementary 3D model, showing how the variation of antennal orientation and the direction of individual units affect the overall auditory diagram

**Supplementary computer 3D model**. Compiled executable together with the source files in MS Visual Basic 5.

