## Supplementary computer 3D model for "Mapping the auditory space of *Culex pipiens* female mosquito in 3D": readme.txt

Lapshin Vorontsov 2022 Mapping the auditory space of Culex pipiens female mosquito in 3D, supplementary model
Authors:     Dmitry Vorontsov, Dmitry Lapshin


Diagrams.exe           compiled executable
	requires:
	XYZF0_D.DAT    pre-caclulated 3D diagram of a single sensory unit 

	may require:
	Diagrams.INI   settings
	msvbvm50.dll   MS Visual Basic 5 runtime library, required in some newere versions of MS Windows

Source code

Diagrams.vbp         'main project file (opened with MS VB 5)
Diagrams.vbw
CommonDialog.bas     'modules
INIwork.bas
Module1.bas
MainWindow.frm
MainWindow.frx

dia.bas              'calculation of individual 3D diagram (not required for the main project)
